# Constraints on paternal care capacity favor a monogamous mating system in the group-living gobiid fish *Trimma marinae*

**DOI:** 10.1101/2025.09.06.674417

**Authors:** Kota Kambe, Kazuya Fukuda, Tomoki Sunobe

**Author notes:** Correspondence: Kazuya Fukuda). 1-15-1 Kitazato, Minami-ku, Sagamihara, Kanagawa 252-0373, Japan. Equal contribution (Kota Kambe, Kazuya Fukuda).

## Abstract

We examined why *Trimma marinae* is monogamous despite high potential for polygyny. Previous work suggested males can care for up to three clutches, but four may exceed their capacity. We therefore tested whether a single female can saturate a male’s care capacity. In established pairs, 49 spawning intervals had a median interval of 2 days. Because eggs hatch on day 3, 75.5% of intervals produced clutch overlap. Under these conditions, mating with an additional female would require care of four clutches, likely exceeding male capacity. These findings support female mate guarding as a strategy to monopolize limited paternal care.

## Introduction

Monogamy is a mating system in which males and females typically mate with a single partner, often maintaining this relationship beyond one reproductive cycle; given the difficulty in defining monogamy unambiguously across animal lineages (Kvarnemo 2018; Bales et al. 2021), this study adopts the definition traditionally used in fish behavioral ecology, following Barlow (1988) and Whiteman and Côté (2004). The ecological conditions under which monogamy is evolutionarily stable have long been investigated (e.g., Emlen and Oring 1977; Shuster and Wade 2003). Even when focusing on marine fish, at least six ecological factors have been proposed to facilitate the evolution of monogamy: biparental care, habitat limitations, low population density/low mate availability/low mobility, increased reproductive efficiency, territorial defense, and the net benefit of sequestering a single mate (Whiteman and Côté 2004). Because there is a great diversity of marine fish, accumulating knowledge of the ecological features of monogamous species is expected to strengthen existing theories and generate new hypotheses.

*Trimma marinae* is a small (approx. 25 mm TL), group-living gobiid fish found on coral reef slopes in enclosed bays of the Western Pacific (Shibukawa 2004). Although information on the ecology of *T. marinae* in the wild is limited, at least at the sampling sites of Fukuda et al. (2017a) and the present study, this species forms aggregations around isolated dead coral patches on muddy bottoms. These aggregations consist of multiple sexually mature males and females. Based on aquarium observations by Fukuda et al. (2017a), it is presumed that reproduction occurs among individuals within the aggregated groups. Additionally, depending on the season, immature juveniles that have settled after the pelagic larval phase also join the aggregations (Fukuda et al. 2017a). The framework of Emlen and Oring (1977) predicts that species in environments with a high potential for acquiring additional and monopolizable mates will exhibit a high environmental potential for polygamy. Based on this framework, *T. marinae* is predicted to favor the evolution of a polygamous mating system because it inhabits groups where potential mates are spatially clustered. However, contrary to this prediction, Fukuda et al. (2017a) revealed that *T. marinae* has a monogamous mating system. According to Fukuda et al. (2017a), *T. marinae* tends to establish continuous reproductive pairs, and most spawning occurs within these pairs. This monogamous mating system primarily results from female mate guarding (i.e., female–female competition). Males do not guard females, but defend the spawning nest from other males and protect the eggs from predation by other group members. At 25°C, eggs hatch three days after spawning. Paired females often deposit the next clutch in the same nest before the previous clutch has hatched. Consequently, males occasionally care for multiple clutches laid by the same female simultaneously. Fukuda et al. (2017a) observed that males in established pairs exhibit reduced aggression toward unpaired females when there are no eggs in their nests and occasionally mate with additional females. These observations suggest that, although males are intrinsically motivated to seek extra-pair mating opportunities, these opportunities are restricted by female mate guarding. This perspective also explains why males are constrained to mating with a single female. In contrast to males, Fukuda et al. (2017a) suggested that females exhibit behaviors facilitating monogamy (i.e., mate guarding) because the limited paternal care capacity of males renders this mating system adaptive for females. By preventing her partner from mating with other females, a female should be able to avoid sharing the male’s limited parental investment, thereby increasing her reproductive success. This hypothesis is supported by occasional observations from Fukuda et al. (2017a) showing that while males could care for up to three clutches, they failed to care for all of them successfully when simultaneously guarding four clutches, with some eggs decomposing or being cannibalized. These observations suggest that the paternal care capacity of *T. marinae* males is constrained. However, that study did not quantitatively investigate whether a male’s care capacity is already saturated by the reproductive output of a single female partner during continuous reproduction within typical monogamous pairs.

To address this question, the present study investigated the spawning interval of continuous pairs under laboratory conditions. Our aim was to determine the number of clutches a male must typically care for simultaneously, and thereby examine the hypothesis that females obtain benefits from mate guarding.

## Materials and methods

The data used in this study were obtained from a rearing observation previously reported by Fukuda et al. (2017a). The materials and methods for specimen collection and rearing conditions, which are identical to the original report, are briefly summarized below. A single entire social group consisting of 41 individuals (14 males and 27 females) was collected by SCUBA at Amami Oshima, Kagoshima Prefecture, Japan, and transported to the laboratory. As 16 individuals (6 males and 10 females) died during the collection and transport process, the remaining 25 individuals (8 males and 17 females) were subcutaneously tagged for individual identification and housed in a 120 × 45 × 45 cm tank at 25°C. Females were significantly larger than males in both the collected (14 males, mean ± SE = 24.5 ± 0.17 mm TL, range = 23.0–25.0 mm; 27 females, 25.8 ± 0.19 mm TL, range = 24.0–27.0 mm; Welch’s *t*-test, *t* = −5.32, *P* < 0.01) and experimental groups (8 males, 25.0 ± 0.18 mm TL, range = 23.5–25.0 mm; 17 females, 26.0 ± 0.22 mm TL, range = 24.0–27.0 mm; Mann–Whitney *U*-test, *U* = 50, *P* < 0.01). Although the number of individuals decreased, the sex ratio in our experimental setup did not significantly differ from that of the original wild group (Fisher’s exact test, *P* = 0.76). Therefore, we consider it likely that the reproductive behavior and mating system observed in the experimental tank reflect those occurring in the wild. Ten polyvinyl chloride pipes were halved as spawning nests. Reproductive behavior was observed between 04:00 and 12:00 for 37 consecutive days. At the rearing temperature of 25°C, the eggs of *T. marinae* hatch on the third day post-fertilization, with hatching triggered by it becoming dark (sunset). Fukuda et al. (2017a) reported no significant correlation in body size between males and females forming continuous pairs (Spearman’s rank correlation coefficient, *r* = −0.39, *P* > 0.05, n = 21).

Following the criteria of Fukuda et al. (2017a), a male–female pair was considered an established monogamous bond if pairing behavior was observed for more than 2 days. As *T. marinae* typically reproduces in monogamous pairs, we recorded spawning intervals exclusively for females in these established pairs. To clarify representative spawning intervals, we analyzed only females that spawned at least twice and recorded the number of days between their successive spawning events.

## Results

A total of 49 spawning intervals were recorded from established pairs, with a median interval of 2 days; this means that females typically spawned approximately 48 hours after the previous spawning (IQR = 1–3 days). As eggs of *T. marinae* hatch on the third day post-fertilization, this reproductive pace indicates that males may be required to care for two clutches simultaneously. The full frequency distribution of the spawning intervals, which ranged from 1 to 13 days, is shown in Fig. 1. Of the 49 total recorded intervals, 37 (75.5%) were 3 days or less (i.e., intervals of 1, 2, or 3 days); this reproductive pace requires males to care for at least two clutches simultaneously. Similarly, 26 of these intervals (53.1%) were 2 days or less (requiring care for three or more clutches), and 12 (24.5%) were just 1 day (requiring care for four clutches).

**Fig 1.**
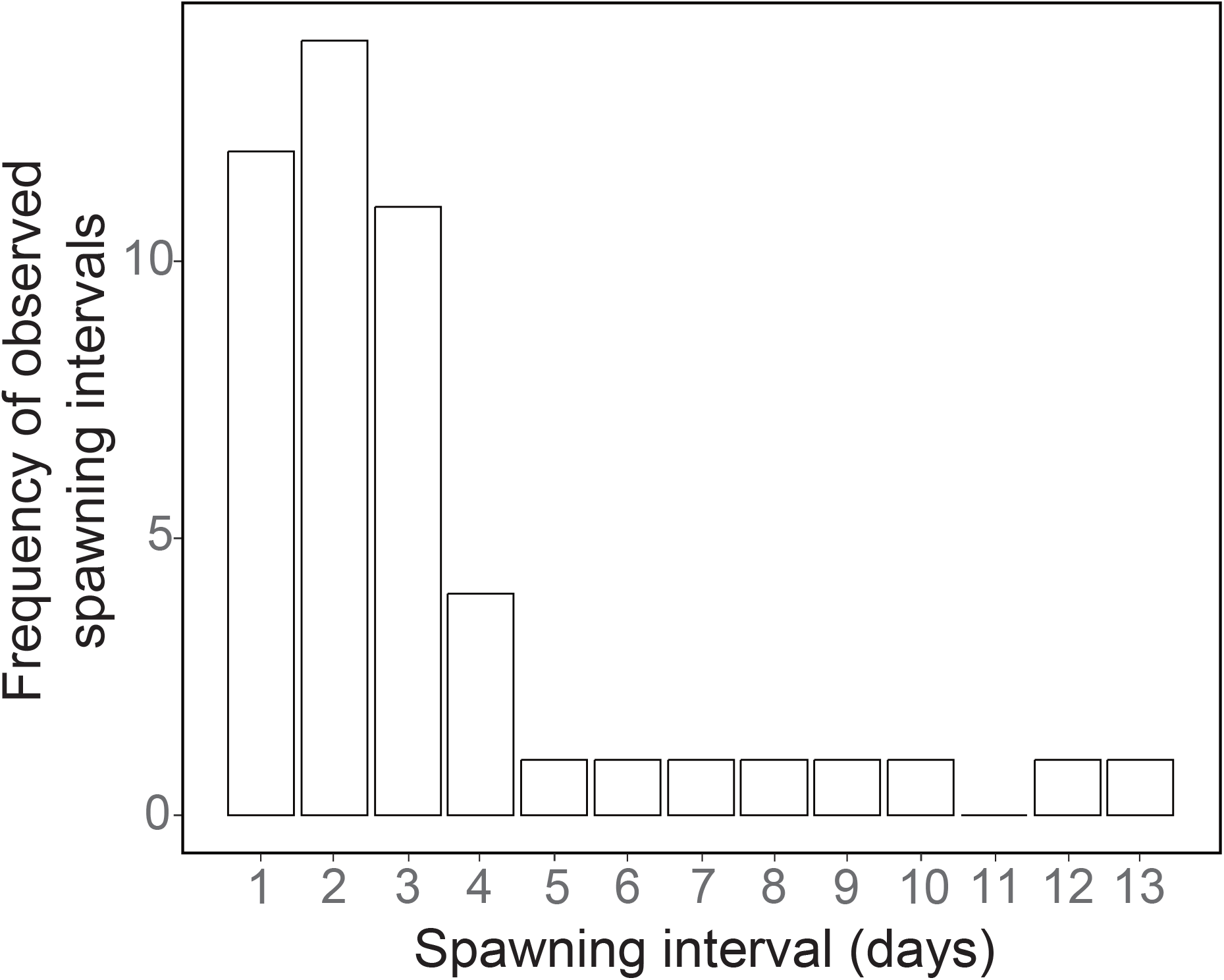
Histogram of the spawning intervals distribution (days). The x-axis indicates the number of days between spawning events within monogamous pairs

## Discussion

This study demonstrated that in 75.5% of reproductive events within monogamous *T. marinae* pairs, females spawn their next clutch before the previous one hatches. This indicates that males routinely face opportunities to care for two clutches simultaneously during a typical reproductive cycle, even when paired with a single partner. This situation suggests that mating with a second female would create a high probability of the male being required to care for four clutches, a number exceeding the male’s paternal care capacity. Furthermore, in 53.1% of reproductive events, the spawning pace required males to care for three clutches simultaneously, indicating that a single female’s reproductive cycle is already sufficient to saturate a male’s paternal care capacity. Therefore, to ensure that their own eggs receive sufficient care until hatching, it would be adaptive for a female to prevent her partner from acquiring additional mates. This female behavior may explain the evolution of a monogamous mating system in *T. marinae*, despite their high environmental potential for polygamy. The present findings offer support for the hypothesis proposed by Fukuda et al. (2017a). Since Fukuda et al. (2017a) and the present study examined only a single wild group, it remains unclear whether the female-biased sex ratio is a consistent feature across all populations or seasons in this species. However, if this skewed sex ratio represents the conditions in the wild, at least during the reproductive season, resulting female-skewed operational sex ratio (OSR), may have led to conditions that favored the evolution of female mate guarding and subsequent monogamy in *T. marinae*. Indeed, in the monogamous pipefish, *Corythoichthys haematopterus*, which also exhibits a female-biased sex ratio, female mate guarding is suggested to contribute to the maintenance of monogamy (Matsumoto and Yanagisawa 2001). This behavior is considered to be promoted by the skewed OSR (Sogabe and Yanagisawa 2007). Further field studies examining the sex ratio across multiple groups and seasons are required to assess the validity of the hypothesis discussed in this study.

While potential ultimate drivers that explain the evolution of monogamy are diverse (e.g., Whiteman and Côté 2004; Klug 2018), the framework proposed by Whiteman and Côté (2004) provides an effective basis for explaining the evolutionary background of monogamy in marine fishes, including *T. marinae*. Among the drivers proposed by this framework, "biparental care" is ruled out for this species because *T. marinae* exhibits paternal care. Similarly, the formation of aggregated social groups where potential mates are abundant contradicts hypotheses based on "habitat limitations" or "low population density/low mate availability/low mobility." The "territorial defense" hypothesis is also unlikely because there is no evidence of cooperative territorial defense in this species. Regarding "increased reproductive efficiency," we cannot currently evaluate its potential contribution, as data on the relationship between pair duration and reproductive efficiency are unavailable for this species. On the other hand, the "net benefit of sequestering a single mate" hypothesis appears to align well with our findings. Specifically, the female strategy of monopolizing a male’s limited parental care capacity corresponds to this driver.

Whiteman and Côté (2004) applied this hypothesis to explain the evolutionary driver of monogamy in *Oxymonacanthus longirostris*, which exhibits monogamy driven by female mate guarding. Therefore, it is plausible to hypothesize that the saturation of male paternal care capacity, resulting from the short spawning intervals of females demonstrated in this study, serves as a potential ultimate driver for the evolution of monogamy in *T. marinae*.

One possible factor contributing to the limited paternal care capacity of *T. marinae* is that males are smaller than females (Fukuda et al. 2017a). This pattern contrasts with that of the polygynous *Trimma* species (e.g., *T. okinawae* [Sunobe and Nakazono 1990; Manabe et al. 2007], *T. grammistes* [Fukuda et al. 2017b], *T. caudomaculatum* [Tomatsu et al. 2018], *T. emeryi* and *T. hayashii* [Fukuda and Sunobe 2020]) (Table 1). In these species, the larger male body size likely enables the care of multiple clutches from several females simultaneously, thus leaving females with little need to compete for paternal care. Indeed, in *T. caudomaculatum*, it has been observed that three females spawned simultaneously in a single male’s nest without competition (Tomatsu et al. 2018). In *T. okinawae, T. grammistes, T. emeryi*, and *T. hayashii*, the largest individual in a harem is confirmed or presumed to reproduce as a male, and in *T. caudomaculatum*, a few large males in the social group monopolize mating opportunities (Table 1). Consequently, females in these polygynous species almost always mate with males larger than themselves. On the other hand, *T. marinae* shows no significant correlation in body size between males and females forming continuous pairs (Fukuda et al. 2017a). Although this suggests that paired males are not always smaller than their partners, given that females are significantly larger than males at the population level, females are likely to pair with males that are similar in size or smaller than themselves. In *T. marinae*, therefore, the restrictedmale body size relative to females likely constrains their capacity to care for the clutches produced by the relatively larger females. This constraint is thought to drive intrasexual competition among females to monopolize a male’s parental care, favoring the evolution of monogamy. The case of *T. taylori*, the only other *Trimma* species reported with a similar sexual size dimorphism to *T. marinae*, lends further support to this hypothesis; it is also suggested to be monogamous due to female competition over limited paternal care (Oyama et al. 2023). This pattern suggests that a small male body size relative to females has been a potential factor in the evolution of monogamy, at least within the *Trimma* lineage. Notably, within the *Trimma* lineage, the evolution of monogamy as observed in *T. marinae* and *T. taylori* does not appear to be closely associated with habitat characteristics (Table 1). This suggests that the evolution of monogamy in this lineage may have been influenced more strongly by constraints on paternal care capacity than by habitat-related factors. However, because field observations of mating systems are still limited in *Trimma*, further accumulation of field data will be necessary to identify the ecological conditions that favored the evolution of monogamy in this lineage. Cases suggesting that constraints on paternal care induce female mate guarding, ultimately leading to the evolution of monogamous mating systems, have been previously reported. For example, in the coral-dwelling goby *Paragobiodon xanthosomus*, female intrasexual competition driven by limitations in paternal care is suggested to be a key factor in the establishment of its monogamous mating system (Wong et al. 2008). This process is similar to the one proposed for *T. marinae*. Given that parental care is commonly provided only by males in gobiid fishes, further studies may identify additional species exhibiting monogamy due to similar selective pressures, even in environments with a high potential for polygyny.

**Table 1.** Summary of mating systems, group structures, and body sizes of seven *Trimma* species reported in previous studies. Data sources: *Trimma marinae* (Fukuda et al. 2017a; Senou et al. 2021); *Trimma taylori* (Senou et al. 2021; Oyama et al. 2023); *Trimma okinawae* (Sunobe and Nakazono 1990; Manabe et al. 2007; Senou et al. 2021); *Trimma grammistes* (Fukuda et al. 2017b; Senou et al. 2021); *Trimma caudomaculatum* (Saeki et al. 2005; Tomatsu et al. 2018; Senou et al. 2021); *Trimma emeryi* (Fukuda and Sunobe 2020; Senou et al. 2021) and *Trimma hayashii* (Hagiwara and Winterbottom 2007; Fukuda and Sunobe 2020; Senou et al. 2021). *Group* refers to specific social groups analyzed in each study, while *Total* refers to pooled data for the study individuals. *Body length* is presented as total length, except for *T. okinawae*, which is presented as standard length. In the *Size Relationship* column, *M > F* and *F > M* indicate that males are significantly larger than females and vice versa, respectively, while *NS* indicates no significant difference.

| Species | Mating system & Group structure | Male body length (mean ± SE) | Female body length (mean ± SE) | Size Relationship | Habitat characteristics |
| --- | --- | --- | --- | --- | --- |
| *Trimma marinae* | Monogamy | Group A: 24.5 ± 0.6 mm TL, n = 14 | Group A: 25.8 ± 1.0 mm TL, n = 27 | Group A: M < F | Overview: Aggregation, hovering<br>Depth: 15–26m<br>Habitat: Drop-offs (sloping or vertical, with caves) and isolated dead corals |
| *Trimma taylori* | Monogamy | Group A: 27.9 ± 1.0 mm TL, n = 9<br>Group B: 28.1 ± 1.1 mm TL, n = 9<br>Group C: 28.5 ± 0.9 mm TL, n = 14<br>Group D: 28.7 ± 1.2 mm TL, n = 16 | Group A: 28.9 ± 1.0 mm TL, n = 17<br>Group B: 29.0 ± 1.3 mm TL, n = 15<br>Group C: 28.9 ± 0.9 mm TL, n = 49<br>Group D: 29.9 ± 0.9 mm TL, n = 26 | Group A, D: M < F<br>Group B, C: NS<br>(Females are generally larger than males) | Overview: Aggregation, hovering<br>Depth: 20–44m<br>Habitat: Rocky caves |
| *Trimma okinawae* | Harem polygyny | Total: 25.4 ± 1.6 mm SL, n = 63 | Total: 22.5 ± 1.8 mm SL, n = 146 | Total: M > F | Overview: Solitary/small group (harem), cryptobenthic<br>Depth: 4–7m<br>Habitat: Cave ceilings, rock sloping, holes, and the underside of table corals |
| *Trimma grammistes* | Harem polygyny | Total: 29.7 ± 4.3 mm TL, n = 22 | Total: 29.6 ± 2.6 mm TL, n = 45 | Total: NS<br>(Males tend to be larger than females in a social group) | Overview: Solitary/small group (harem), cryptobenthic<br>Depth: 18–24m<br>Habitat: Rocky reefs |
| *Trimma caudomaculatum* | Polygyny in multi-male group | Group A: 34.5 ± 1.0 mm TL, n = 4<br>Group B: 32.5 ± 0.5 mm TL, n = 3<br>Group C: 32.5 ± 0.5 mm TL, n = 7<br>Group D: 34.0 ± 1.5 mm TL, n = 12 | Group A: 31.0 ± 2.0 mm TL, n = 9<br>Group B: 29.5 ± 1.0 mm TL, n = 14<br>Group C: 29.2 ± 1.4 mm TL, n = 35<br>Group D: 29.5 ± 1.5 mm TL, n = 100 | Group A, B, C, D: M > F | Overview: Aggregation, hovering<br>Depth: 1–5m<br>Habitat: Patch reefs with overhanging holes or tunnel structures |
| *Trimma emeryi* | Harem polygyny (putative) | Group A: 22 mm TL, n = 1<br>Group B: 22.5 mm TL, n = 1<br>Group C: 23.5 mm TL, n = 1 | Group A: 21.0 ± 0.5 mm TL, n = 2<br>Group B: 21 ± 0 mm TL, n = 2<br>Group C: 20.8 ± 1.8 mm TL, n = 2 | Total: M > F | Overview: Solitary/small group (harem), cryptobenthic<br>Depth: 6–23m<br>Habitat: Deep off with cave and crevices |
| *Trimma hayashii* | Harem polygyny (putative) | Group A: 26.5 mm TL, n = 1<br>Group B: 27 mm TL, n = 1<br>Group C: 24.5 mm TL, n = 1<br>Group D: 25 mm TL, n = 1<br>Group E: 23.5 mm TL, n = 1<br>Group F: 24.5 mm TL, n = 1<br>Group G: 26.5 mm TL, n = 1 | Group A: 24.0 ± 0.6 mm TL, n = 3<br>Group B: 24.0 ± 0.5 mm TL, n = 2<br>Group C: 22.8 ± 0.3 mm TL, n = 2<br>Group D: 21.0 mm TL, n = 1<br>Group E: 22.5 ± 0.3 mm TL, n = 3<br>Group F: 23.0 ± 1.0 mm TL, n = 2<br>Group G: 24 mm TL, n = 1 | Total: M > F | Overview: Solitary/small group (harem), cryptobenthic<br>Depth: 6–20m<br>Habitat: Drop off with cave and crevice, reef slopes with corals |

## Acknowledgments

We are grateful to the members of the Laboratory of Fish Behavioral Ecology, Tokyo University of Marine Science and Technology for their advice and support during our study. Thanks are also due to S. Yokoyama for their assistance during the study.

## Author contributions

Conceptualization, Methodology: KF, TS; Investigation: KF; Formal analysis: KK; Writing - original draft preparation: KK; Writing - review and editing: KF, TS; Funding acquisition: TS; Resources: KF, TS; Supervision: TS.

## Data availability

The datasets are available from the corresponding author on reasonable request.

## Ethical approval

At Tokyo University of Marine Science and Technology, there are no institutional ethical regulations for fish. Therefore, all procedures adhered to the ASAB/ABS guidelines for the treatment of animals in behavioral research.

